# Molecular diagnosis of new isolate of tomato yellow leaf curl virus in Iraq

**DOI:** 10.1101/2021.04.02.438091

**Authors:** Ali Alabde, Osamah Alisawi, Fadhal Al Fadhal

## Abstract

Tomato yield and quality in Iraq have been threatened by a variable range of infections caused by tomato yellow leaf curl virus. In previous studies, the TYLCV isolates were partially characterized using molecular tests for small fragments not the entire length of the virus. Sample of TYLCV-infected tomato has applied in this study to diagnose complete sequence of TYLCV isolate. Three sets of primers that belong to three well-identified strains in Iraq were used in a PCR technique and interestingly the results were negative. A new Iraqi isolate has been characterized as a first novel Iraqi isolate detected ever using next generation (NGS) and bioinformatics techniques. The NGS platform has produced about 78,232,062 paired reads of the TYLCV-infected tomato var. Oula F1. The complete raw reads of the infected variety have been analyzed using RepeatExplorer pipeline and Map to reference. The full sequence of TYLCV was reconstructed and extracted to consist of 2770 nt and then deposited in Genbank under accession number MT583814. Copy numbers and genome proportion of this sequence have been calculated that were 3523 and 0,086% respectively. Phylogenetic analyses of full nucleotide sequences confirmed close relationship to Iranian isolate (TYLCV-Kahnooj) than other published viruses. Additionally, a and β DNA satellites have not discovered in the TYLCV-infected sample.

## Introduction

The family Geminiviridae is recently divided into seven genera: *Becurtovirus*, *Begomovirus*, *Curtovirus*, *Eragrovirus*, *Mastrevirus*, *Topocuvirus* and *Tur ncurtovirus.* Host range, insect vector, genome organization and genome-wide pairwise sequence identities have been applied to classify this family (Varsani *et al.*, 2017). Tomato yellow leaf curl virus (TYLCV) belongs to genus *Begomovirus* and has a monopartite single-stranded DNA (ssDNA) encapsidated in geminate particles. TYLCV is transmitted by the whitefly, *Bemisia tabaci* (Genn.) that effectively spread virus particles through persistent manner (Panno *et al.*, 2016). The disease caused by this virus is the most important disease on tomato in Iraq and worldwide (Mansoor *et al.*, 2003; Salati *et al.,* 2002; Al-Kuwaiti *et al.*, 2013). Infected plants become stunted with chlorotic leaves, curled-up margins, and abscised flowers. Since the first infection was recorded in Israel 1939-1940, multiple regions have been identified to have isolates from this virus over the world and classified to different viral species (Antignus and Cohen 1994; Delatte *et al.*, 2004; Pico *et al.*, 1996; Polston and Anderson 1997). Tomato yellow leaf curl Sardinia virus TYLCSV infected tomatoes in Israel and Sardinia, and was the first sequenced isolate in 1991(Kheyr-Pour *et al.*, 1991; Navot *et al.*, 1991), and since that date many isolates have been sequenced from various locations (Moriones and Navas-Castillo 2000). In Iraq, the disease was detected based on symptoms assay and serological tests. Recently, most popular molecular works on this virus have been achieved by PCR based on amplified short fragments of TYLCV genome such as coat protein domain (Al-Ani *et al.*, 2010; Reddy *et al.*, 2005; Maruthi *et al.*, 2007). Further, many tomato genotypes have been screened against TYLCV isolates to characterize similarities and differences through these genotypes (Al Tamimi, 2020). TYLCV has been isolated not only from tomato but also from various plants such as eggplant, melon, cucumber, sesame, cowpea in addition to weeds as a primary source of infection like *Malva parviflora*, *Chenopodium murale* and *Chenopodium quinoa* (Al-Jubouri *et al.*, 2014). Al-Kuwaiti *et al* (2013) have reported complete sequence of TYLCV that showed 99% identity to Spain strain (Acc. AJ489258) confirming its existence in Iraq. In this study, we utilized next generation sequencing and bioinformatics techniques for the first time in Iraq to find out new isolate with complete sequence of TYLCV. Furthermore, phylogeny analyses have been performed to understand how these isolates related to each other.

## Materials and methods

### Plant material

From open fields of Najaf governorate, the main region of tomato production in Iraq, fresh and young leaves of TYLCV-infected tomato variety Oula F1 (Seminis Seeds Company, USA) were harvested. This variety is considered the most popular and TYLCV-sensitive commercial variety of tomato in the middle of Iraq.

### DNA isolation, PCR and NGS sequencing

To extract the DNA, the young leaves (about 2 gm) of the infected tomato were applied using cetyl-trimethylammonium bromide (CTAB) method with minor modifications (Doyle and Doyle 1990). Three sets of primers that belong to strains of TYLCV-IS, TYLCV-ES and TYLCV-Mld were applied in a PCR technique to check their infection within examined sample (Table 1) (Anfoka *et al.*, 2005).

**Table 1.**
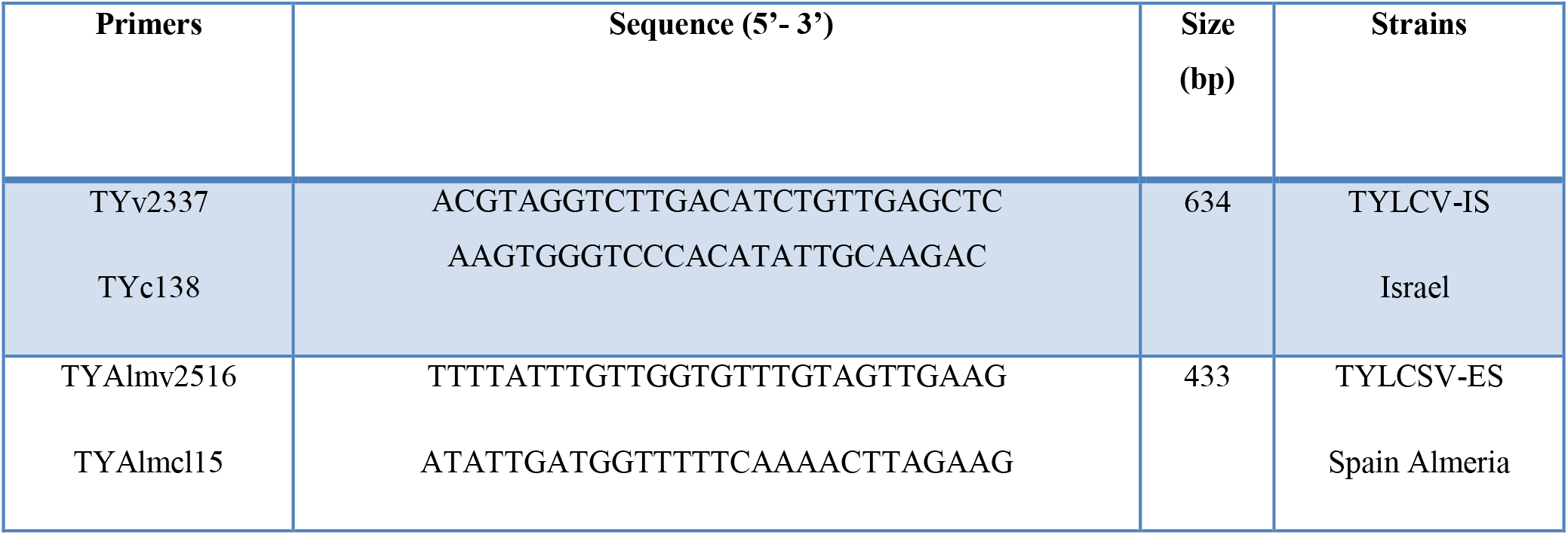
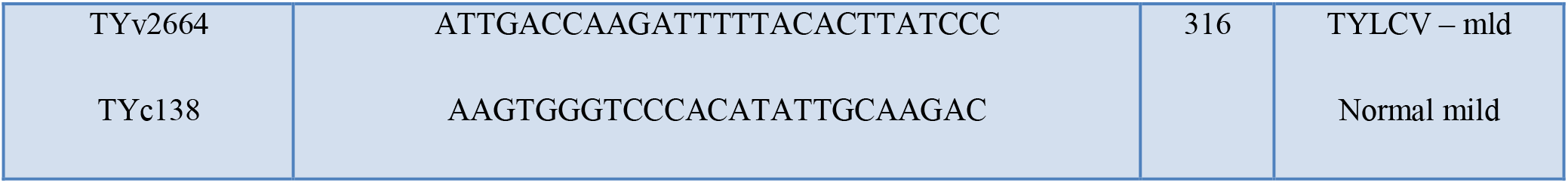
Three primers of main strains of TYLCV in Iraq that used to amplify and characterize virus sequence in the examined sample.

PCR conditions were an initial denaturation of 10 min at 95°C, 49 cycles of 30 s at 95°C, 20 s at 55-63°C and 30 s at 72°C followed by a final extension of 30 s at 72°C. To obtain the whole genome sequence, the extracted genomic DNA sample was sent to sequencing at DNA-link Company, Republic of Korea. The genome was sequenced by using the Novaseq6000 2×150bp reads technique and application WGS (PCR Free550) based on the manufacturer’s procedure.

### RepeatExplorer program

To collect virus sequences from the whole data (NGS) of infected tomato, RepeatExplorer pipeline was applied to recognize these kinds of sequences in forms of clusters within repetitive DNA blocks (Novák *et al.* 2013). The targeted clusters of RepeatExplorer were extracted, and then Repbase dataset (Jurka *et al.* 2005), and Basic Local Alignment Search Tool (Altschul *et al.* 1990) were used to identify contigs of each cluster. Furthermore, the extracted sequences were mapped to well-identified plant virus sequences from DPVweb (Adams and Antoniw 2005).

### Map to reference

Using Geneious program V. 11 (Kearse *et al.* 2012), the raw reads were assembled against TYLCV Iraq (Acc. JQ354991.1) as a reference sequence that diagnosed by Al-Kuwaiti *et al.* (2013) and then consensus sequence was collected. Mapping to reference was repeated against consensus sequence (TYLCV) to confirm and obtain the actual number of assembled reads. Also, the raw reads were applied to assemble against a and β DNA satellites. The outcomes showed in a report that has a number of assembled reads and total used reads in addition to the frequent overlapped reads. This data used to calculate copy numbers (Number of assembled reads x read length/reference sequence length) and genome proportions (Number of assembled reads / numbers of total NGS reads x 100) (Mustafa *et al.* 2018). Finally, the two consensus sequences from RepeatExplorer and Map to reference were aligned using pairwise alignment in Geneious program to collect the right length of TYLCV isolate.

### Phylogenetic analysis

The maximum likelihood (ML) method was applied to find phylogeny model using MEGA7 (Tamura *et al.* 2013). The Geneious program V. 11 (Kearse *et al.* 2012) was used for alignment and optimized manually. The entire nucleotide sequences of 13 main TYLCV isolates including our isolate of interest, Kahnooj and Iraq isolates were applied to phylogeny study. ClustalW alignment was used for extracting sequences about 2770 nt for each. The phylogenetic tree reconstructed using General Time Reversible (GTR). Bayesian phylogeny inference was used for analysis with Bayesian inference of phylogeny (MrBayes 3.2.6) (Huelsenbeck and Ronquist 2001).

## Results

The extracted DNA concentration and quality were 418 ng\ml and 1.92 which was in a high quality level for sequencing approach (Fig. 1. A). PCR has surprisingly failed to get amplified sequences for the three TYLCV isolates indicating that our sample has probably a new virus sequence. High quality raw reads have been obtained from NGS Illumina platform with about 78,232,062 paired reads; 150 bp read length and 36.9% GC content (Fig. 1. B).

**Fig. 1.**
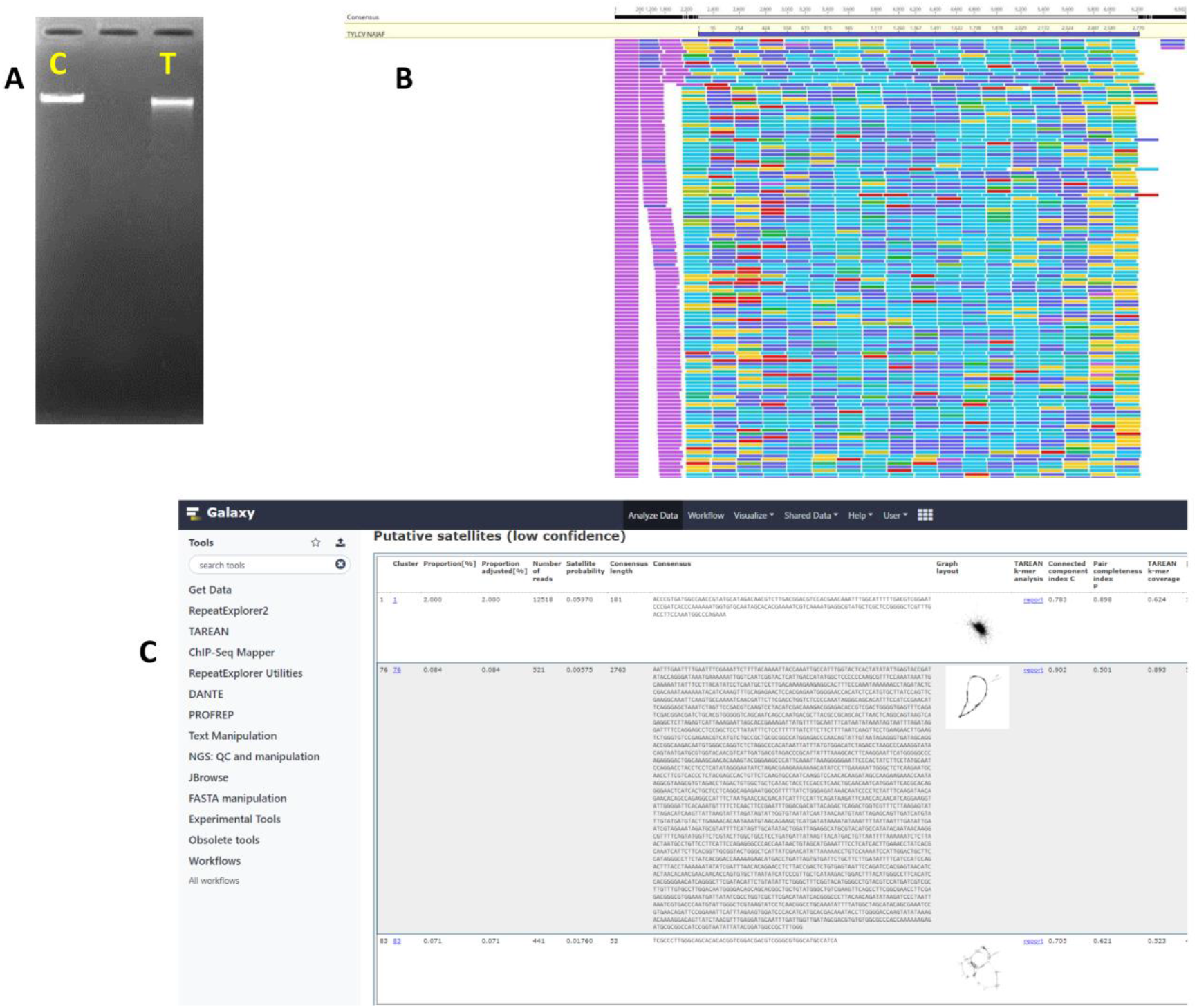
A: Gel electrophoresis of genomic DNA of TYLCV-infected tomato var. Oula F1 (T), in comparison with a control genomic DNA (C). B: Raw reads mapped to complete sequence of TYLCV-Najaf isolate (2770 nt), the assembled reads were 65,303 that used to calculate genome proportion and copy numbers. C: RepeatExplorer report showed cluster number 76 for TYLCV sequence (2763 nt) that picked up as a putative satellite due to tandemly arranged feature within raw reads.

RepeatExplorer has been clustered the whole uploaded raw reads that showed tandemly repeated sequences, TYLCV sequences involved in cluster number 76 as a putative satellite with a genome proportion 0,084% and the consensus sequence was 2763 nt (Fig. 1. C). On the other side, the report of Map to reference revealed that 65,303 reads have been assembled to count genome proportion and copy numbers of the isolate that were 0,086% and 3523 respectively whereas the consensus sequence was 2770 nt. In order to obtain complete sequence, the aligned consensus sequences from both RepeatExplorer and Map to reference showed 100% similarity and the longest sequence was extracted. Lastly, we determined 2770 nt of TYLCV and named as Najaf isolate and then deposited in Genbank under accession number MT583814.1.

Phylogenetic analyses showed what we have proven earlier by pairwise alignment between Najaf isolate and each of Kahnooj and Iraq isolates. Kahnooj was most likely similar to Najaf isolate than Iraq. TYLCV nucleotide and amino acid sequence alignments showed that Najaf isolate is closely related to Kahnooj isolate the Iranian TYLCV in 93.8% pairwise identity while it was 90.3% with Iraq isolate (Fig. 2). However, polymorphic sequences have shown higher rate in intergenic region (IR) (75%), V2 (91.9%), C1 (Rep) (93.1%), and C4 (88.3%) than other coding regions V1 (97.3%), C3 (95.9%) and C2 (96.5%).

**Fig. 2.**
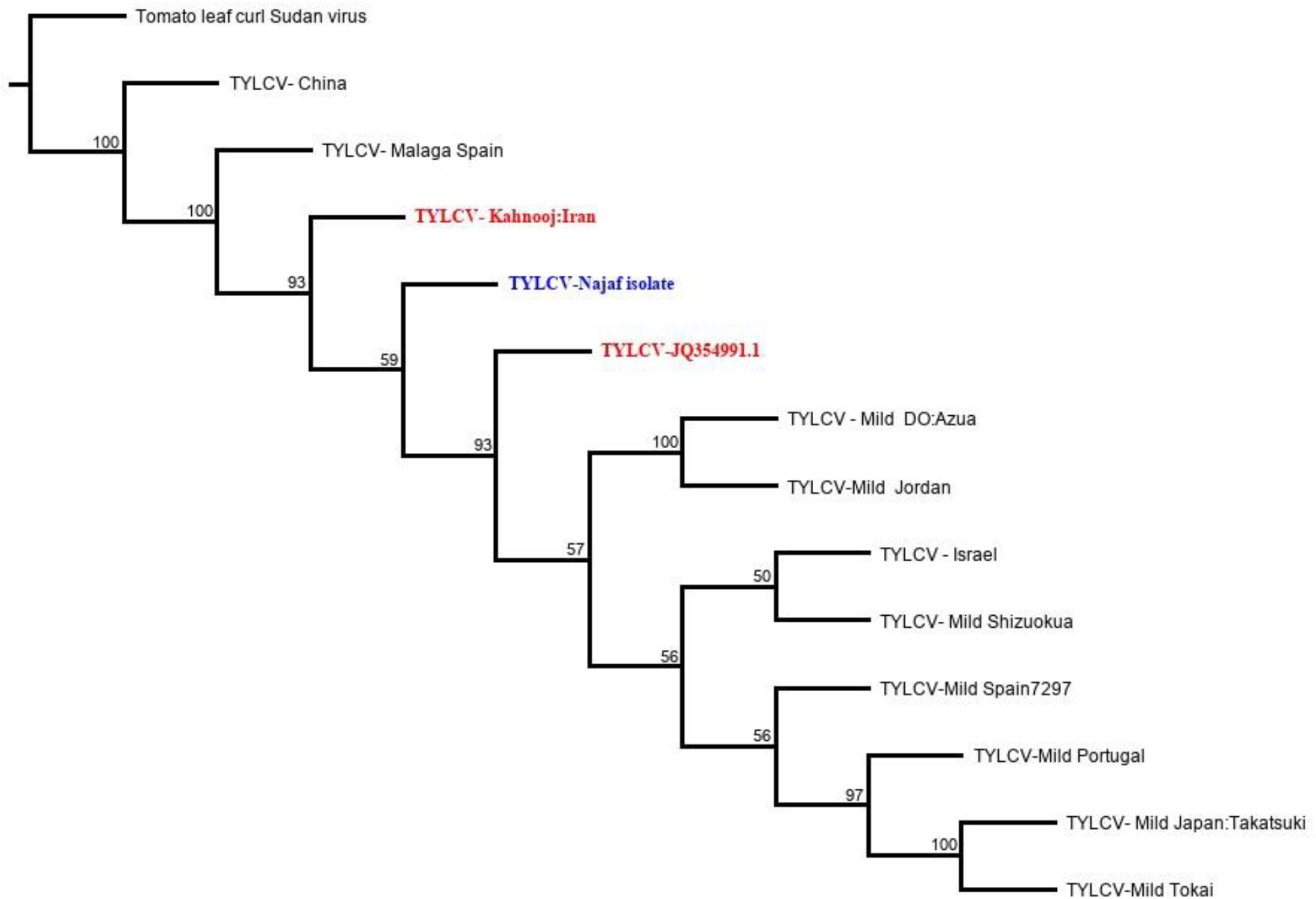
Phylogenetic tree of 13 full lengths TYLCV isolates showing close relationship between Najaf and Kahnooj isolates. The tree was generated using Geneious tree builder with bootstrape analyses and 50% support, the out group member was *Tomato leaf curl Sudan virus*.

Our research has been extended trying to find DNA-a and β satellites using Map to reference tool but we did not get any positive hits.

## Discussion

The deep sequencing of whole genome sequencing technique has revolutionized genomic research collecting a massive data of the host genome with complete sequences of cellular components. Additionally, this method is considered as unbiased approach to detect plant viruses with no previous knowledge about the suspected elements. The NGS technique does not use specific primers, antibodies, and virus-specific reagents to diagnose pathogenic viruses (Adams *et al.*, 2009). In this method, full image of plant viruses and complex infection could be investigated in the host genome unlikely to biased methods that focus on almost one expected virus (Jones *et al.*, 2017). In Iraq, most of molecular tests concentrated on partial sequences of TYLCV such as coat protein domain to detect the virus and apply these amplified parts in multiple alignments and phylogeny analyses. The only full-length sequenced of this virus from Iraq was Iraq isolate (JQ354991.1) that achieved by Al-Kuwaiti *et al* (2013) and named as TYLCV Iraq but it shared 99% similarity to Spain strain (Acc. AJ489258), Reunion Islands (Acc. AM409201) and Mauritius (Acc. HM448447). Additionally, Sadeq Al-Kuwaiti (2013) has applied rolling circle amplification (RCA) and non-degenerate primers to catch the full length of virus. Therefore, our new isolate is considered the first full sequenced Iraqi isolate of TYLCV with totally new data yet. We found Kahnooj isolate from Iran is closer than other isolates even from Iraq with 93.8 % similarity. It is highly expected that Najaf and Kahnooj isolates are sharing the same ancestor and most likely the virus was imported into Iraq from Iran by international trading (Mali *et al.*, 2003; Gibbs *et al.*, 2008 Al-Kuwaiti 2013).

Mapping reads to the reference has revealed the estimated copy numbers of this virus was 3523 within infected cell. Copy number value has been always considered as a good sign for variation and variability of endogenous components within host genome especially transposable elements including viruses (Mustafa *et al.* 2018; Alisawi 2019). Interestingly, this is one of NGS and bioinformatics benefits to calculate copy numbers of the components of interest and this approach has been applied in many studies to predict viral copies (Yang *et al.*, 2017; Alisawi 2019). So far, the number of copies indicates virus particles indeed infected the host genome. In fact, this little amount of copies strongly explains the early stage of symptoms emerging in the infected sample. However, further works need to be achieved regarding isolate distribution and diversity over tomato varieties and different regions in Iraq.

We could not reveal the existence of DNA satellites in our infected sample probably due to moderate level of infection in the examined sample. A and β DNA satellites are often associated with begomoviruses in their replication, movement within infected plant and horizontal transmission between plants, as well as involved in induction of virus symptoms and suppression of transcriptional gene silencing. This result is consistent with most published outcomes that reported the importance of DNA satellites for disease severity and inducing symptoms. However, we do expect to find these satellites in severely TYLCV-infected plants as no more researches have been covered DNA satellites in Iraq (Laufs *et al.*, 1995; Saunders *et al.*, 2004; Cui *et al.*, 2005; Yang *et al.*, 2011; Amin *et al.*, 2011; Zhou, 2013).

## Conclusion

This work concluded that whole genome sequencing technique alongside bioinformatics is highly recommended to detect complete sequence of TYLCV isolate within infected tomato genome raw reads. Consequently, a clear insight of virus pathway and identity has been revealed by phylogenetic analysis. More investigations need to be done for detecting full sequences of highly pathogenic plant viruses in Iraq.

## Statement of Ethics

The authors have no ethical conflicts to disclose.

## Author Contributions

A.A. carried out the experiment. O.A. analyzed the NGS data and wrote the manuscript with support from F.A. F.A. helped supervise the study.

